# A monotone single index model for missing-at-random longitudinal proportion data

**DOI:** 10.1101/2022.01.20.477170

**Authors:** Satwik Acharyya, Debdeep Pati, Dipankar Bandyopadhyay, Shumei Sun

## Abstract

Beta distributions are commonly used to model proportion valued response variables, commonly encountered in longitudinal studies. In this article, we develop semi-parametric Beta regression models for proportion valued responses, where the aggregate covariate effect is summarized and flexibly modeled, using a interpretable monotone time-varying single index transform of a linear combination of the potential covariates. We utilize the potential of single index models, which are effective dimension reduction tools and accommodate link function misspecification in generalized linear mixed models. Our Bayesian methodology incorporates the missing-at-random feature of the proportion response, and utilize Hamiltonian Monte Carlo sampling to conduct inference. We explore finite-sample frequentist properties of our estimates, and assess the robustness via detailed simulation studies. Finally, we illustrate our methodology via application to a motivating longitudinal dataset on obesity research recording proportion body fat.

## 1 INTRODUCTION

Research in various biomedical disciplines and public health generates data, where the primary response variables are constrained in a compact interval, say in (0,1), rather than the whole real line. For example, consider our motivating Fels longitudinal study (Roche 1992, FLS) recording proportion body fat (pbf), a popular clinical biomarker in obesity research,for study subjects at longitudinal time-points, along with various important covariates, such as gender, age, body mass index, etc. Figure 1a presents the raw density histogram of the response ‘pbf’ ∈ (0,1), packed across all subjects and time-points. For conducting regression, one can potentially transform the response variable to the real line, and use conventional approaches (Qiu, Song, & Tan 2008). However, such vanilla approaches pose both modeling and computational issues, given that inference can be sensitive to the transformations used, and parameters in the transformed scale rarely carry similar interpretation as in the original model. In a quest to conduct direct modeling of such responses, the Beta density (Gupta & Nadarajah 2004), a continuous log-concave density, is often the density of choice due to its versatility in accommodating a variety of unimodal shapes on a compact interval, and thereby address non-Gaussianity, and data skewness(Smithson & Verkuilen 2006). Under a generalized linear mixed model (GLMM) framework, a reparametrized beta density (and associated regression) assists us to conveniently connect the model covariates to the proportion response (Ferrari & Cribari-Neto 2004), with a subject-specific random effects in clustered/longitudinal studies (Hunger, Döring, & Holle 2012; Petterle, Bonat, & Scarpin 2019). Other popular distributions modeling proportion responses include the beta rectangular (Bayes, Bazán, & García 2012; Hahn 2008), simplex (Barndorff-Nielsen & Jørgensen 1991), logistic normal (Aitchison 1986), and the Bessel (Barreto-Souza, Mayrink, & Simas 2020).

**FIGURE 1.**
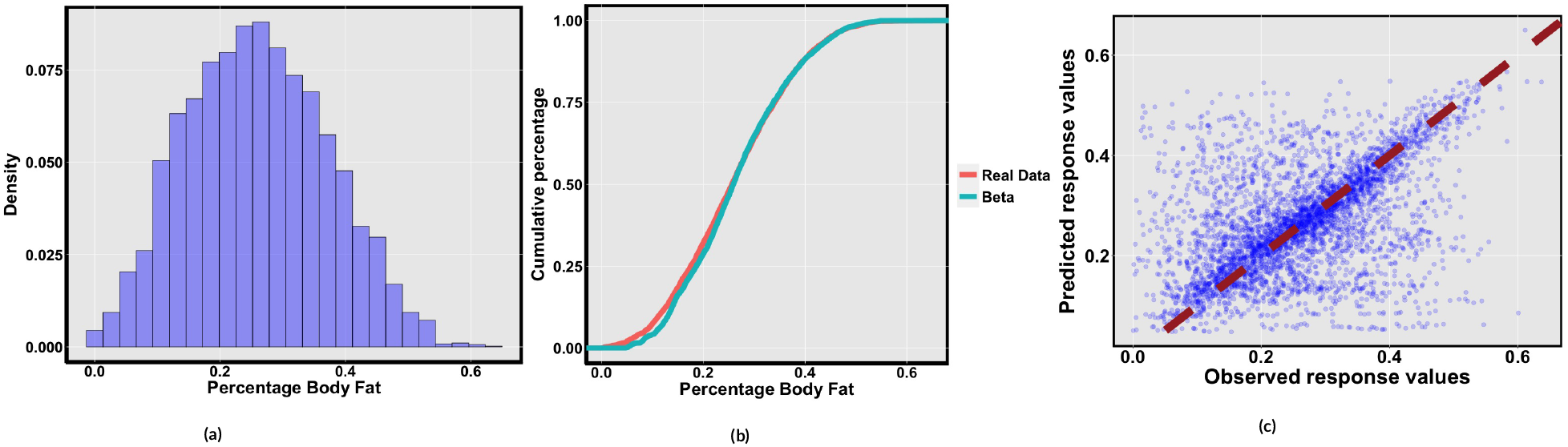
*The left panel shows the histogram of percentage body fat (pbf). The middle panel provides empirically calculated cumulative distribution of the response variable i.e. percentage body fat and the same after fitting beta regression combined with monotone single index (2). The right panel plots the predicted vs observed response variables from BR-MSIM* (*μ*_it_, *ψ*) *model*.

Although linear regression (LR) are simple and commonly used procedures for evaluating covariate-response relationships, they are inadequate for inference and prediction under violations of the LR assumptions. In biomedical (obesity) research, index measures, such as the Charlson comorbidity index (Afolabi et al. 2020), combines information from an array of observed characteristics into a *single value,* thereby providing important unobserved traits of a subject (Wu & Tu 2016). However, a majority of these indices were developed on empirical grounds, and lacks sound statistical justification. On the other hand, a single-index model (Stoker 1986, SIM) provides a simple, interpretable framework for quantifying a complex, possibly non-linear relationship between a response Y_i_ and the p > 1 dimensional covariate vector X_i_ = (X_i1_,…, X_ip_), where the conditional expectation of Y_i_ |X_i_ can be expressed as an unknown, univariate function g(.) of the scalar index 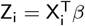, where *β* = (*β*_1_,…, *β_p_*) is an unknown index vector (more details in Section 2). This SIM specification accommodates both non-linear main effects, and higher order interactions (determined by the function g(.)), and thus offer a pragmatic compromise between a fully parametric LR, and other nonparametric formulation (Dhara, Lipsitz, Pati, & Sinha 2020). However, (clinical) interpretation can be compromised(Foster, Taylor, & Nan 2013), if the shape of g(.) is left completely unspecified. Hence, monotone single-index models (Balabdaoui, Durot, & Jankowski 2019; Groeneboom & Hendrickx 2019) have evolved, leading to straightforward clinical interpretation, and ease of inference. In this paper, we model the longitudinal proportion response using a BR, where the logit transform of its mean is flexibly modeled via a monotone single index transform of a linear combination of time-varying covariates. The justification behind enforcing monotonicity of the index function appears in the next section. Under a Bayesian paradigm powered by the Hamiltonian Monte Carlo (MMC) sampling (Betancourt, Byrne, Livingstone, & Girolami 2017), a missing-at-random (MAR) assumption accommodates seamless handling of the missing responses within the Bayesian updating scheme.

The rest of the article proceeds as follows: After a brief introduction to BR and the challenges associated with a SIM, Section 2 presents the monotone SIM specification via Bernstein polynomials (BP), and the consequent adaptation to handle MAR missingness. In Section 3, we develop the Bayesian estimation scheme, with prior specifications, posterior inference via HMC, and associated Bayesian model selection tools. Application to the motivating FELS data, with relevant posterior summaries and prediction accuracy appear in Section 4. In Section 5, we use synthetic data to evaluate the finite sample properties, and robustness of our proposal, in light of existing alternatives. Finally, some concluding comments and future developments appear in Section 6.

## 2 STATISTICAL MODEL

### 2.1 Single-index Beta Regression

Let y_it_ is the observed proportion response ∈ (0,1), and X_it_ ∈ ℝ^p^ the corresponding predictors at tth (t = 1,…, T_i_) time point for the ith (i = 1,…, n) subject. We model y_it_, conditional on the covariates as

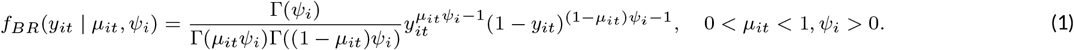

where, *μ*_it_ = E(y_it_) is the mean parameter, with individual specific precision parameter *ψ*_i_.This is denoted as y_it_ ~ Beta(*μ*_it_*ψ*_i_, (1 – *μ*_it_)*ψ*_i_). Now, to propose the beta regression, the covariates are connected to the mean *μ*_it_ as 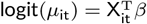, where *β* ∈ ℝ^p^ isthe vector of regression coefficients. The variance component *ψ*_it_ is left unspecified (to be estimated using some priors as in our case), or estimated via assigning a link function, such as log, to the covariates.

Over the years, there has been considerable effort in constructing more flexible mean functions of multivariate covariates. In this context, the SIMs(Hardle, Hall, & Ichimura 1993), where logit(*μ*_it_) is expressed as 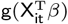, is typically viewed as a bridge between a multiple linear regression, and a non-parametric regression problem. The SIM do not suffer from the curse of dimensionality due to the reduction to an univariate index variable from the multidimensional predictor set (Yu & Ruppert 2002). This dimension reduction property enhances computational scalability, and preserves the flexibility of the model through utilization of non-linear functions. They are also advantageous in the context of misspecification of the non-linear link function on 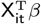. The function g(·) is called a link function, and the p-variate coefficient vector *β* is called the index vector. The linear combination 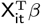 is referred as the index for the predictor X_it_. In a SIM, one aims to estimate both the link function g and the index vector *β* simultaneously. The problem of estimating g is the same as a non-parametric univariate regression problem. An advantage of using a SIM over a usual multiple linear regression problem is that once can achieve a higher predictive power as we are considering a function of a linear combination of the covariates (Carroll, Fan, Gijbels, & Wand 1997). SIM is also a special case of projection pursuit regression (PPR) (Friedman & Stuetzle 1981) with a single component, thereby providing simpler interpretation over the multi-component PPR. Thus, in addition to better prediction performance, the SIM provides appealing interpretation of the covariate effects on the response.

Despite its popularity, the SIM brings in certain challenges for statistical inference. The parameters (g, *β*) are not jointly identified. Constraints on (g, *β*),such as monotonicity on g (Foster et al. 2013), and a unit norm constraint on *β* facilitate identifiability and the interpretation of the covariate effect on the response. With g(.) assumed monotone, there are additional advantages in interpretation and usefulness of the index 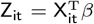 for subject i. Without loss of generality, if g is monotone non-decreasing, the expected value of the response will increase or remain the same with increase in Z_it_, and a prespecified threshold on 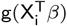 enjoys a one-on-one correspondence to an equivalent threshold on 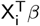, allowing elegant interpretation. Moreover, if the coefficient/parameter corresponding to a specific X_it_ is positive, then increasing the value of X_it_ (with other covariates remaining fixed) will result in a higher value of the index, thus increasing the expected value of the response. Based on these interpretations, clinicians may aim to lower the expected adverse response by taking appropriate steps to lower the index, Z_it_, by implementing a plan todecrease/in-crease a particular X_it_. These physical interpretations of g and β are not always available when the link function g is unconstrained. In the following, we present the details of our monotone SIM for longitudinal proportion data, and the adjustments to handle missingness.

### 2.2 Monotone SIM

Classical methods (Ayer, Brunk, Ewing, Reid, & Silverman 1955; Brunk 1955) of imposing monotonicity requires constrained optimization to ensure the monotonicity and smoothness of the link function. Isotonic regression i.e, fitting a monotone function g to data points (y_i_, x_i_) involves finding a weighted least-squares fit p ∈ ℝ^n^ to the vector y ∈ ℝ^n^, with weights vector w ∈ ℝ^n^ subject to a set of constraints of the kind p_i_ ≤ p_i+1_.Then, the isotonic regression problem corresponds to the following quadratic program: 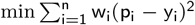 subject to p_i_ ≤ p_i+1_.This can be solved using a simple iterative algorithm called the pool adjacent violators algorithm (De Leeuw, Hornik, & Mair 2010). In presence of the single index, such optimization routines can be problematic.

To circumvent this issue, we use a smooth BP basis (Farouki 2012) to model the link function. In the following, we describe how using a BP basis reduces the problem of estimating a monotone link function to a constrained linear regression problem. In a Bayesian context, optimization is avoided by placing suitable prior distributions on the constrained space, and then sampling from the posterior distribution to produce estimates that satisfy the constraints. The SIM connects the linear predictors with a non-linear function to model the logit linked mean of the response variable as

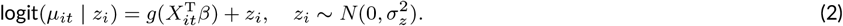

where g(.) is a unknown monotone link function on ℝ → ℝ, and z_i_ denote subject-specific random effects. We impose a standard restriction on *β* i.e. || *β*|| = 1, to ensure identifiability of the model. Detailed discussion appears in Subsection 2.3. This assumption helps to provide a better interpretation of the response variable i.e., pbf, wrt. the time component. We model the monotone link function g(·) with BP and the monotonicity is ensured through imposing necessary restriction on the coefficients of the polynomial. The BP of degree M is defined as

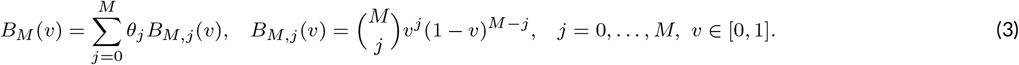

where, B_M_(v) is non-decreasing if the coefficients of the polynomial are non-decreasing i.e. *θ*_0_ < *θ*_2_ < … < *θ*_M_ (Chak, Madras, & Smith 2005). Following Souris, Bhattacharya, and Pati (2018), we scale the input variable 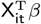 because the domain of BP is defined on [0,1] interval. Using Cauchy-Schwarz inequality and identifiability constraints, we have 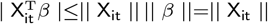 where || *β*|| = 1. We consider *c* = max || X_it_||, and transform 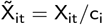, leading to the identity 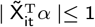. Another transformation is required on the BP B_M,j_(v) = p_j_(u)/(M + 1), v ∈ [0,1] to provide support on [–1,1], where, p_j_(v) is a Beta(j + 1, M — j + 1) density. We take the transformation W = 2V – 1, where p_j_(v) is the density of the variable V and the density of W is q_j_(w) = p_j_{(w + 1)/2}/2 for w ∈ [—1,1].The transformed BP basis for j = 0,…, M is then defined as

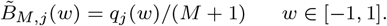

The monotone SIM with transformed BP basis is given as

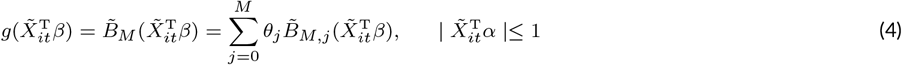

where 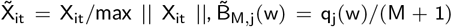 and q_j_(w) is a transformed beta density with | w |< 1. Now, 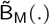 is non-decreasing because of the order-restriction on the basis coefficients 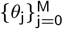. We define an equivalent transformation on 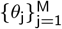 and set *ϕ*_0_ = *θ*_0_, *ϕ*_1_ = *θ*_1_ – *θ*_0_, …, *ϕ*_M_ = *θ*_M_ – *θ*_M-1_, such that *ϕ*_k_ ≥ 0 for k = 1,…, M. We write AΦ = *θ* where *θ* = [*θ*_0_, *θ*_1_,…, *θ*_M_], Φ = [*ϕ*_0_, *ϕ*_1_,…, *ϕ*_M_] and A is a (M + 1) × (M + 1) dimensional matrix with all the lower triangle and diagonal entries are 1.Thus, (2) can be rewritten as

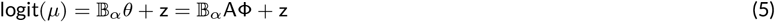

where 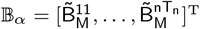 is a nT × (M + 1) matrix with 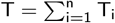 and 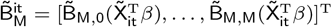.

### 2.3 Identifiability of parameters

Lin and Kulasekera (2007) proved the identifiability constraints for the SIM, under the assumption of non-constant and continuous g, || *β*|| = 1, with the first non-zero element being positive. In our case, the monotonicity property of the function g assists in relaxing the assumption of the first non-zero element of *β* to be positive (Balabdaoui et al. 2019). The proof of the identifiability constraint is also extended to left or right continuous functions instead of continuous functions (Balabdaoui et al. 2019) under i.i.d. setup. Observe that due to the presence of random effects, the well-studied iid setup has been violated. However, this does not pose a significant problem to show identifiability as we shall see below.

To define the model (2), we need at least one observation from a subject at one time point. We define the support of g as 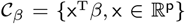. We assume that mean of the response variable exists and (2) holds true for some *γ* = (g, *β*) such that 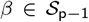 (a unit sphere of dimension p) and 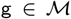, the class of monotone functions on ℝ. We enforce identifiability of the parameters through imposing the following constraints: (a) g(.) is a monotone non-decreasing function i.e., 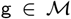, and (b) || *β*|| = 1 i.e. 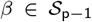. To demonstrate identifiability, we assume that there exists parameters *γ*_1_ = (*g*_1_, *β*_1_) and 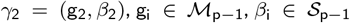, such that f(y_11_, x_11_| γ_1_) = f(y_11_, x_11_| γ_2_). Taking expectation of (2) and applying the inverse logit transformation, we have g_1_ (x^T^*β*_1_) = g_2_(x^T^*β*_2_). To uniquely identify the parameters, it is sufficient to show g_1_(x^T^*β*_1_) = g_2_ (x^T^*β*_2_), which only holds if *β*_1_ = *β*_2_, g_1_ ≡ g_2_. Our final claim holds true by Proposition 5.1 from Balabdaoui et al. (2019).

### 2.4 Handling Missingness

Missing data or incomplete information is a common issue in medical studies. Several techniques to handle missing data have been studied over last few decades using approaches such as data imputation (Harel & Zhou 2007; Rubin 2004; Zhang 2003) and fully Bayes (Daniels & Hogan 2008; Ibrahim, Chen, Lipsitz, & Herring 2005). In absence of a proper clinical justification for missing-not-at-random assumption, we assume the response pbf to be missing at random (MAR), also referred to as ignorable missingness (Little & Rubin 2019). This implies that the missing data mechanism is not dependent on the missing response values.

It is an established fact that unbiased estimates can be obtained from the observed likelihood instead of the joint likelihood of observed and missing data (Seaman, Galati, Jackson, & Carlin 2013). We denote R_it_ as the indicator for the missing response variables at tth time point for the ith individual i.e. R_it_ = 1 if Y_it_ is observed, 0 otherwise. The conditional distribution of missing data mechanism f(R | Y, λ) is identified through the parameter vector λ which is independent from our parameter of interest Θ. In case of MAR, the conditional distribution is independent of the choice of missing response values. Following the definition from Rubin (1976), MAR holds if

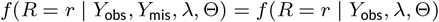

where Y_obs_ and Y_mis_ are observed and missing set of responses, along with λ, the parameter vector of the missingness mechanism. The conditional distribution of (Y_obs_, X, R | λ, Θ) is

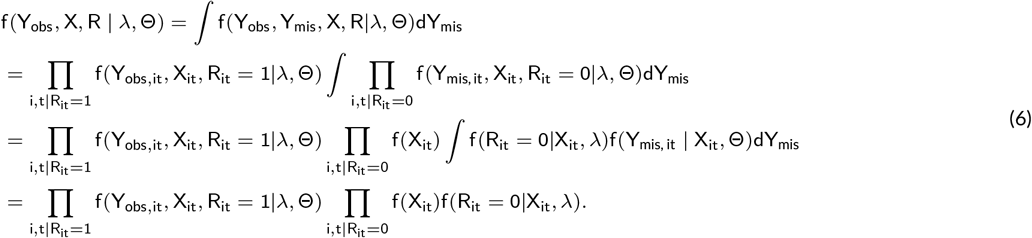

Under MAR assumptions, equation (6) holds, clearly indicating that the inference on Θ (parameter of interest) does not depend on the missing data mechanism. A Bayesian approach can naturally incorporate the uncertainty due to the presence of missing data (Erler et al. 2016). Bayesian methods on missing data can generate posterior samples of the parameters and missing variables from their posterior predictive distribution. In the next section, we outline our hierarchical Bayesian estimation framework that incorporates the missing data model.

## 3 BAYESIAN INFERENCE

### 3.1 Prior specification

Our Bayesian estimation scheme is initiated via specifying the prior distributions on the model parameters. First, we denote *α* = *β*/ || *β* || and place a Gaussian prior on *β*. Next, we posit a standard Gaussian prior on the first coordinate of Φ, while the remaining coordinates together gets a multivariate Gaussian prior N(0, cI_M_) truncated to a positive real line. Anon-informative prior is imposed on the variance of the subject-specific random effects, i.e. 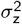 in (2). Following the recommendations from Bandyopadhyay, Galvis, and Lachos (2017), we use a Gamma prior on the precision parameter *ψ* for the BR model in (1).

### 3.2 Posterior Inference

Under the assumptions of independence between the subject-specific random effects and MAR, the observed likelihood of Θ = (*β*, Φ, *ψ, σ*_z_) is

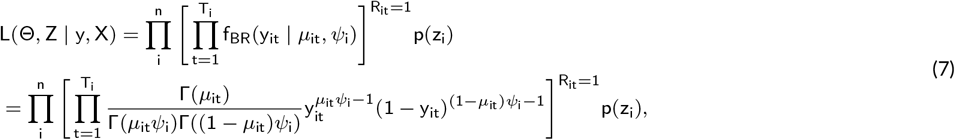

where p(z_i_) denotes the distribution of subject-specific random effects, and *μ*_it_ is presented in (2). The joint posterior distribution can be written as

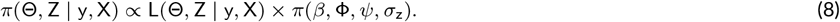

A standard technique to obtain posterior samples is via the implementation of the Markov chain Monte Carlo (MCMC) algorithm, which cycles through the full conditionalsofa parameter given the rest. Instead of using standard MCMC algorithm, we obtain posterior samples by using Hamiltonian Monte Carlo, or HMC (Duane, Kennedy, Pendleton, & Roweth 1987; Neal 1994) and the No-U-turn sampler (NUTS) (Hoffman & Gelman 2014). A probabilistic programming language Stan (Carpenter et al. 2017; Stan Development Team 2019) has been developed for Bayesian inference by combining the HMC and NUTS sampler. Stan is scalable for large datasets, and often achieves faster convergence compared to other available software, such as WinBUGS (Lunn, Spiegelhalter, Thomas, & Best 2009), JAGS (Plummer 2003), and others. We fit our BR model (1) with Stan by specifying the observed likelihood (7) and prior distributions (3.1).

### 3.3 Bayesian model selection and influence diagnostics

To assess model goodness of fit, we compared prediction accuracy through several diagnostic measures (Hoeting, Madigan, Raftery, & Volinsky 1999; Vehtari & Ojanen 2012). Such measures can be either information based, or cross-validatory. The Bayesian information criteria (Schwarz 1978, BIC) penalizes the log-likelihood with number of fitted parameters and sample size. WAIC (Gelman, Hwang, & Vehtari 2014) is also another popular fully Bayes information criteria. Computationally efficient WAIC estimates are obtained from out-of-sample prediction with pointwise log posterior predictive density along with an adjustment of number of effective parameters. Following Gelman et al. (2014), the effective number of parameters is computed as

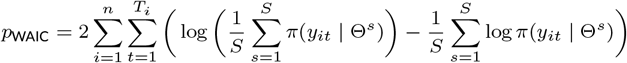

Finally, WAIC is evaluated as

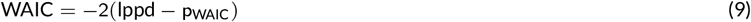

where, lppd is log of pointwise predictive density.

Cross-validation (CV) based criteria are another parallel strategies to capture out-of-sample prediction errors. The diagnostic measures based on CV do not suffer from the problem of over-fitting but they may not be computationally scalable. The conditional predictive ordinate (Dey, Chen, & Chang 1997, CPO) criterion is calculated for an observed point given all other data points. The CPO for the i-th subject at the t-th time point is defined as CPO_it_ = π(y_it_ | y(_it_)) = ∫ π(y_it_ | Θ)π(Θ | y(_it_))dΘ, where y(_it_) denotes the dataset without the observation y_it_. Following Dey et al. (1997), CPO_it_ is numerically computed as

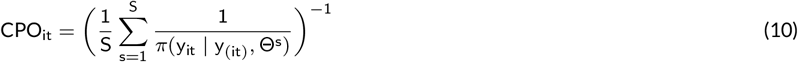

where, S denotes the number of posterior Monte Carlo samples, post-convergence. A higher value of CPO_it_ for a model indicates a better support for the (i, t) th datum. A summary measure is the log pseudo-marginal likelihood (Geisser & Eddy 1979, LPML), defined as

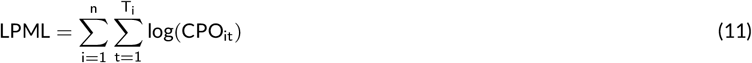

Similar to CPO, a higher value of LPML suggests a better model fit to the data. Under the MAR assumption, we evaluated the model diagnostic measures based only on observed responses (Daniels & Hogan 2008; Ma & Chen 2018).

## 4 APPLICATION: FLS DATA

In this section, we illustrate our BR monotone SIM (BR-MSIM) via application to the motivating FLS data. The FLS (Sun et al. 2007 2008) is the world’s longest and largest longitudinal human growth study that collected the lifetime of repeated measurements on growth, health and body composition of 2,567 European-American participants, as early as 1929. Although participants were enrolled at birth (examined semi-annually until 18 years of age, and biennially thereafter), our current analytical data subset consists of 777 subjects (373 male and 404 females), followed longitudinally since 1976 (year when pbf measurements were included in the study protocol) till 2010. Study subjects exhibit irregular number of time points, with a maximum number of 15 visits. In addition to the response variable (pbf) ∈ (0,1) collected at each time-point, various subject-level covariates, such as gender (Gender, M/F), date of visits (Visit), age (Age, range = 8-83), body mass index (BMI), waist size (Waist), diastolic blood pressure (Dias BP), systolic blood pressure (Sys BP), bicep size(Bicep, in mm), and bio-electrical impedance (BCimped), were also available. Approximately, 17.8% of observations were missing, which we considered MAR. The study was approved by the Institutional Review Boards of the Wright State University and the Virginia Commonwealth University.

We compared the fit of the BR-MSIM to the BR model with linear predictors (BR-Lin), where both models vary with respect to having a subject-specific precision parameter (*ψ*_i_), or an overall precision parameter (*ψ*). The competing models are listed below:

Model 1:y_it_ ~ BR-MSIM(*μ*_it_,*ψ*_i_),
Model 2: y_it_ ~ BR-Lin(*μ*_it_, *ψ*_i_),
Model 3: y_it_ ~ BR-Lin(*μ*_it_, *ψ*),
Model 4:y_it_ ~ BR-MSIM(*μ*_it_, *ψ*).

We choose the optimal value of M ∈ {5,…, 30} for models 1 and 4 based on BIC values. More details about the selection M is deferred to the section S2 of the Supplementary Material. Using the optimal value of M = 22, we compare the four models via WAIC and LPML values (Table 1), calculated using observed data (Ma & Chen 2018). We observe that Model 4 (BR-MSIM, with constant precision parameter *ψ*) has the highest LPML and the lowest WAIC values among the 4 competing models. The BR-MSIM with subject-specific precision *ψ*_i_ is performing poorly because of over-parametrization.

**TABLE 1.** Model comparison with WAIC and LPML values of the 4 models.

|  | Model 1 | Model 2 | Model 3 | Model 4 |
| --- | --- | --- | --- | --- |
| WAIC | -7262.58 | -7341.75 | -12978.72 | -13293.56 |
| LPML | 3595.38 | 3639.56 | 6342.53 | 6490.95 |
| SSE | 16.85 | 17.06 | 7.25 | 6.57 |
| SAE | 211.21 | 214.19 | 129.87 | 122.74 |
| MAPE | 41.88 | 42.64 | 22.03 | 21.05 |
| MAAPE | 0.25 | 0.25 | 0.15 | 0.14 |

Next we report findings from our best-fitted Model 4, i.e., BR-MSIM(*μ*_it_, *ψ*). The estimated posterior mean of regression coefficients with corresponding 95% credible intervals (CI) are presented in Figure 2a. The covariates BMI, Waist, Bicep, and BCimped exhibit significantly higher estimates, compared to the others. All covariates (except Age and Sys BP) have positive regression coefficients.This implies an increase in the estimated single index with an increase in the numerical value for the continuous covariates, or change in category (say, from 0 to 1) for the discrete covariates. However, the covariates Age and Sys BP negatively impact the single index.

**FIGURE 2.**
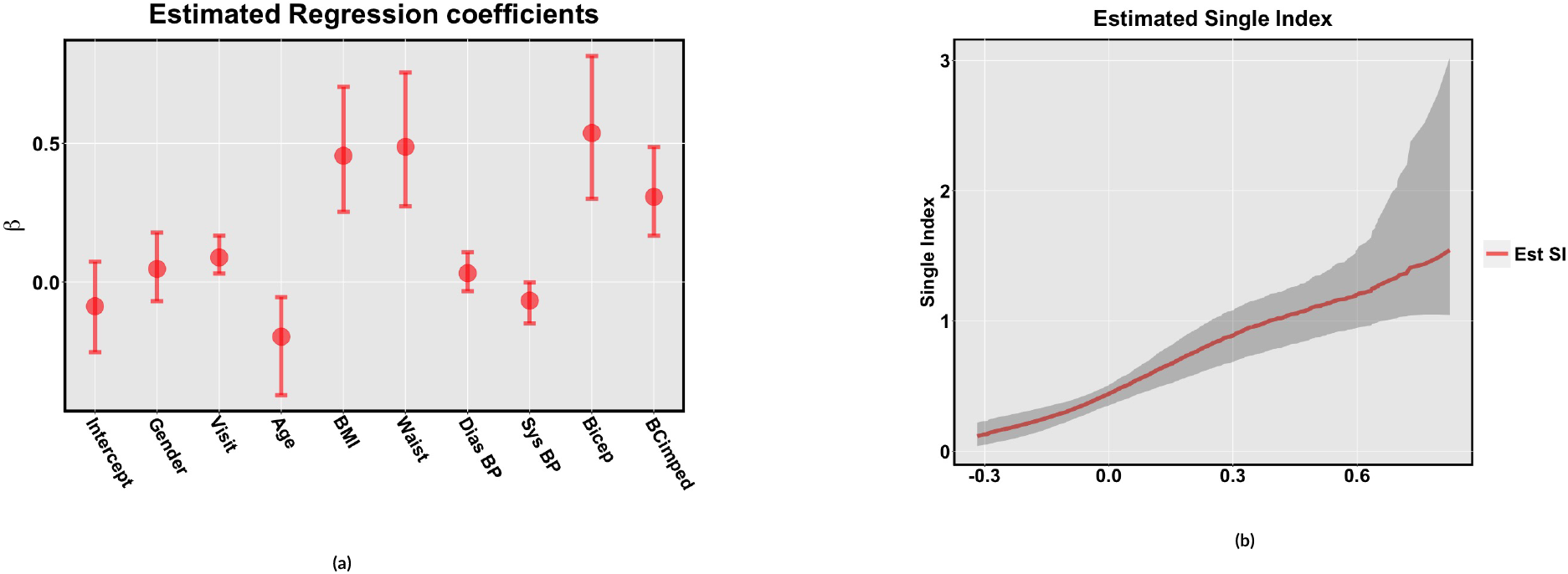
*The estimate of regression coefficients (left panel) and single index (right panel) with 95% credible obtained from BR-MSIM*(*μ*_it_, *ψ*).

Figure 2b plots the estimated single index, with the corresponding 95% CI, denoted by the grey area. To illustrate the monotonicity property of the nonparametric function g(), consider the single index w_it_ for the i-th subject at the t-th time point given by 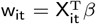, and s_it_ = g(w_it_). Due to this property, we have s_it_1__ > s_it_2__ when w_it_1__ > w_it_2__, where t_1_ and t_2_ are time points of two arbitrary visits of the ith subjects. For an illustration using the FLS data, consider the subject with id = 8 in the FLS data, who is a male with t = 1,…, 7. For this subject, the values of the single index w at the 6th and 7th time-points are w_86_ = 0.18 and w_87_ = 0.30, with the corresponding s_86_ = 0.66 and s_87_ = 0.85, respectively.

Other than WAIC and LPML values, we also used exploratory graphs to assess the goodness of fit. We generate multiple samples from the predictive distribution using the posterior samples. We average over multiple samples to obtain the predicted values of the response variables. Figure 1c represents the association between the predicted values of response variable on the y-axis and observed response variable on the x-axis. The linear trend in the Figure 1c suggests an adequate for the BR-MSIM(*μ*_it_, ψ) model. Next, we calculate prediction accuracy metrics, such as sum of squared errors (SSE), sum of absolute error (SAE), mean absolute percentage error (MAPE), and mean arctangent absolute percentage error (Kim & Kim 2016, MAAPE) to compare the four competing models. These prediction accuracy measures are summarized in Table 1. We observe that Model 4, i.e., the BR-MSIM(*μ*_it_, ψ), has the lowest values corresponding to all metrics, implying superior prediction accuracy compared to the rest three models.

## 5 SIMULATION STUDIES

In this section, we use synthetic data to (a) assess the frequentist finite sample properties (Simulation 1), and (b) assess robustness (Simulation 2), of our proposed monotone SIM.

### 5.1 Simulation 1: Checking frequentist finite sample properties

Here, we investigate theconsistency of single index parametersfor increasing values of n ∈ {100, 200, 300,400, 500}, while setting the percentage of missingness at 20%. To mimic a realistic setting as observed in the FLS electronic health records, varying number of observed (longitudinal) time points are generated via random sampling (with replacement) from {1,…, 10}, with the maximum set to 10. We consider the dimension of regression and basis coefficient to be p = 4 and M = 22 respectively, and fix the true values of the regression coefficients *β* = {2.75,0.85, 2,1.25}, normalized to have unit norm. We set *ψ* = 3, and the basis coefficients *ϕ* are sampled from a vector (0,0.05,0.1,0.2,0.3,0.4), with first entry fixed at 0.15. Posterior estimates were summarized over 50 replicates.

In Figure 3a, we plot the estimated single index function i.e. g(single index) and the associated 95% CIs for n = 100, with the truth overlayed. Under the same setting, we provide the plot for n = 500 in Figure 3b. Furthermore, we split the interval [0,1] with an increment of 0.01, and measured the quantiles of observed and estimated mean (*μ*_it_) at those probabilities. Figures 3c and 3d plot the observed vs estimated quantiles from the mean. To check the consistency of the single index function [g(single index)] parameters, we define a measure of discrepancy as 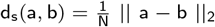, where a, b ∈ ℝ^N^. Figure S3a (Supplementary material) presents the boxplots of 100 × d_s_ between true and estimated single index function for increasing values of n ∈ {100, 200, 300,400, 500} across the 50 replicates. In Figure S3a, the decreasing trend (of Euclidean distances) in the boxplots with increasing n implies consistency of the posterior estimates of single index function parameters. We also report the average bias, MSE and 95% CIs of the regression coefficients *β* in Table 2. All three metrics decreases with increase in the sample size. To ensure convergence of the posterior samples, we also provide several trace plots in figure S1 of the Supplementary Material.

**FIGURE 3.**
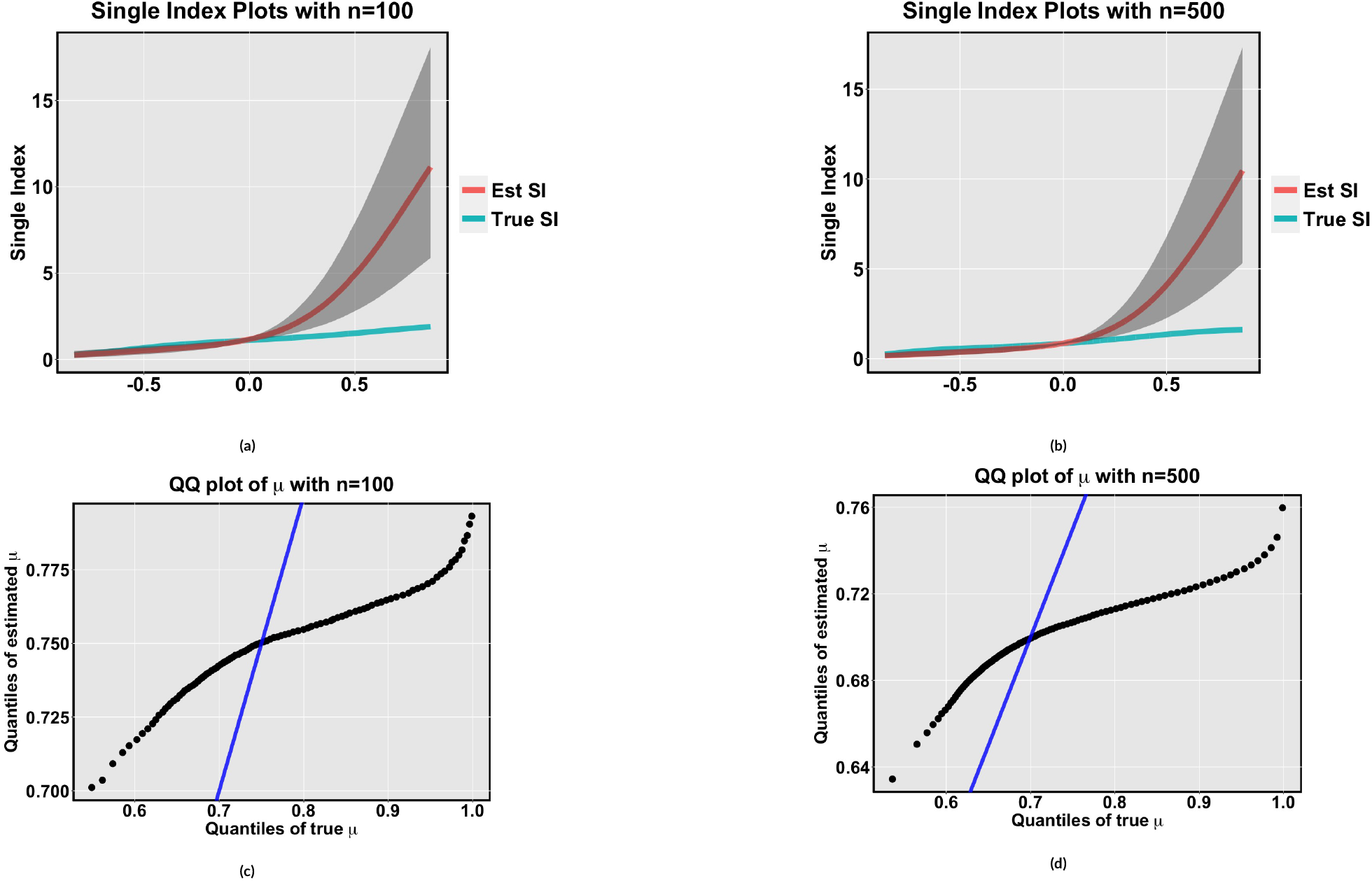
*Figure 3a and Figure 3b are overlaid plots true and estimated single index with* 95% *credible interval for n=100 and* n = 500 *respectively. We provide the quantile plot of observed vs estimated mean* (μ) *with* n = 100 *in Figure 3c and n=500 in Figure 3d.*

**TABLE 2.** *Bias, MSE and 95% credible intervals regression coefficients β obtained from* (2). *All the reported values are averaged over* 50 *replicates.*

|  | n=100 |  |  | n=500 |  |  |
| --- | --- | --- | --- | --- | --- | --- |
|  | Bias | MSE | 95% CI | Bias | MSE | 95% CI |
| $\beta_1$ | 0.24 | 0.42 | (0.01, 0.69) | 0.13 | 0.31 | (0.11, 0.51) |
| $\beta_2$ | -0.27 | 0.34 | (0.01, 0.68) | 0.20 | 0.25 | (0.16, 0.51) |
| $\beta_3$ | -0.04 | 0.40 | (0.02, 0.69) | 0.03 | 0.29 | (0.13, 0.60) |
| $\beta_4$ | -0.16 | 0.38 | (0.01, 0.69) | -0.09 | 0.30 | (0.10, 0.51) |

### 5.2 Simulation 2: Assessing robustness

To assess the robustness of our model, we generate the response variable from a mixture of beta and simplex distribution (Barndorff-Nielsen & Jørgensen 1991), given as (0.8 × Beta + 0.2 × simplex), and fit our proposed BR-MSIM. We present similar plots for this misspecified case, as in Subsection 5.1. Figures 4a and 4b plot the true and estimated single index (with 95% CIs) for n = 100 and n = 500 respectively, when the data is generated from the misspecified data generating distribution. Contrary to Figure S3a, here, we observe an increasing trend in the boxplots of the Euclidean distances (100 × d_s_) with increasing n; see, Figure S3b in the Supplementary Material. The quantile plot of true and estimated mean of the response variable is provided in Figure 4c (n=100) and 4d (n = 500). The effect of misspecification is clear, with the quantiles moving away from the y = x line.

**FIGURE 4.**
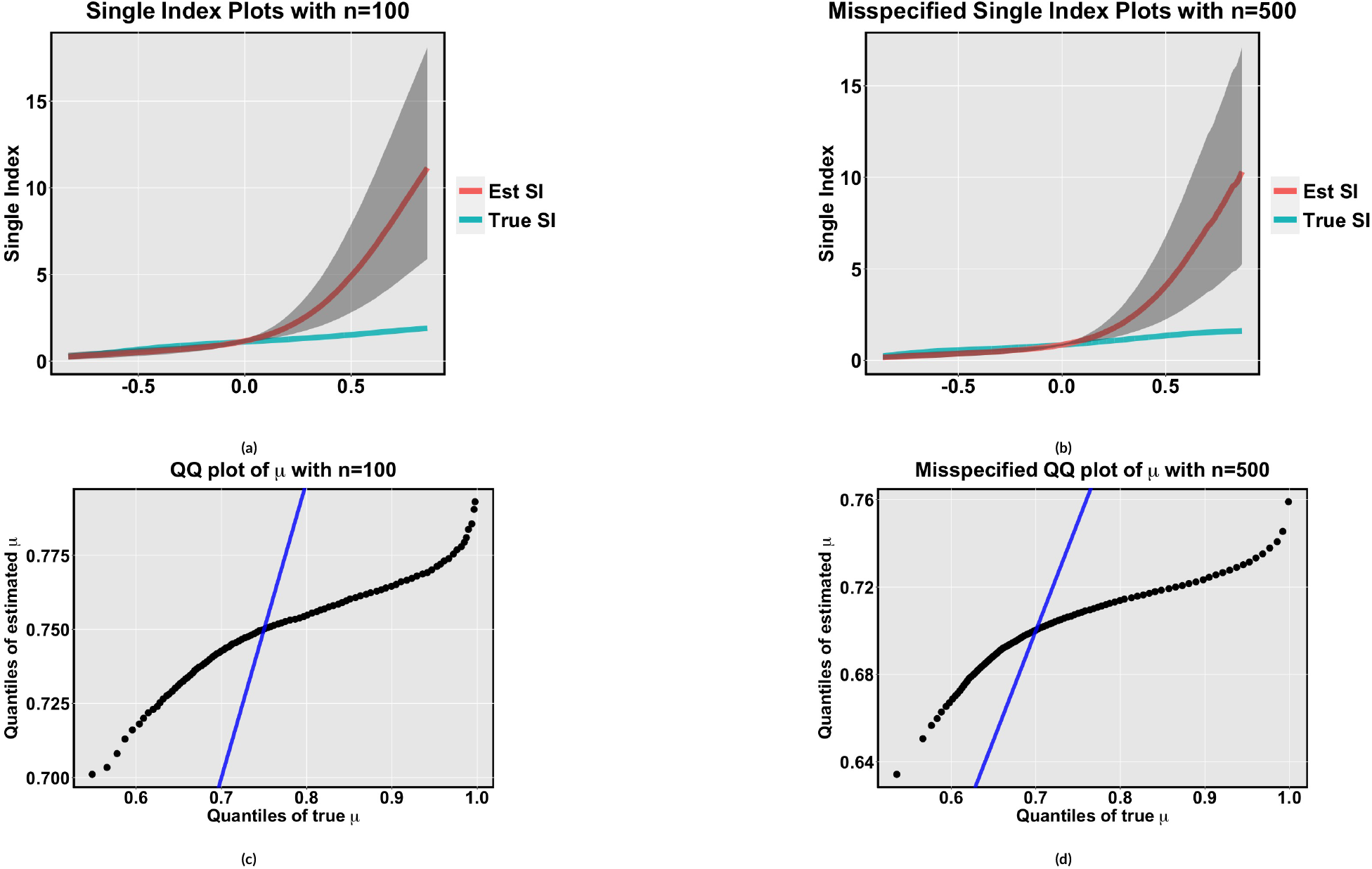
*In Figure 4a and 4b, we provide the plot of single index curves for* n = 100 *and* n = 500 *respectively while the generating data distribution (i.e. a mixture distribution of beta and simplex) is misspecified. Figure 4c and 4d show quantile plot of mean of response variable for* n = 100 *and* n = 500 *respectively in misspecified case.*

## 6 CONCLUSIONS

In this paper, we provide a unique methodology for monotone single index modeling under BR using Bernstein polynomials. This methodology can be extended for any distribution which is supported on the interval (0,1),such as beta rectangular (Bayes et al. 2012; Hahn 2008), simplex (Barndorff-Nielsen & Jørgensen 1991), logistic normal (Aitchison 1986). Our current setup assumes MAR missingness; certainly, this can be extended to MNAR missingness via the popular shared random effects framework (Albert & Follmann 2003) that jointly models the response variable and the (binary) missingness indicator.

## ACKNOWLEDGMENTS

The research was supported by grants R01AG048801, R01DE024984, and P30CA016059 awarded by the United States National Institutes of Health.

## SUPPORTING INFORMATION

The following supporting information is available as part of the online article:

## Notes

### Competing Interest Statement

The authors have declared no competing interest.

