## Supplementary file for "A monotone single index model for missing-at-random longitudinal proportion data"


Satwik Acharya[1], Debdeep Pati[2], Dipankar Bandyopadhyay[3], and Shumei Sun[3]

[1]Department of Biostatistics, University of Michigan, Ann Arbor, MI, USA
[2]Department of Statistics, Texas A&M University, College Station, TX, USA
[3]Department of Biostatistics, Virginia Commonwealth University, Richmond, Virginia, USA


### S1 Diagnostic Check

To investigate the convergence and mixing of the mean parameters, we randomly selected 4 coordinates of $\mu$. In the top row of Figure S1, we show the trace plots of those coordinates with $n = 100$ and the same in bottom row with $n = 500$. Figure S1 suggests good mixing of the parameters for both values of $n$.

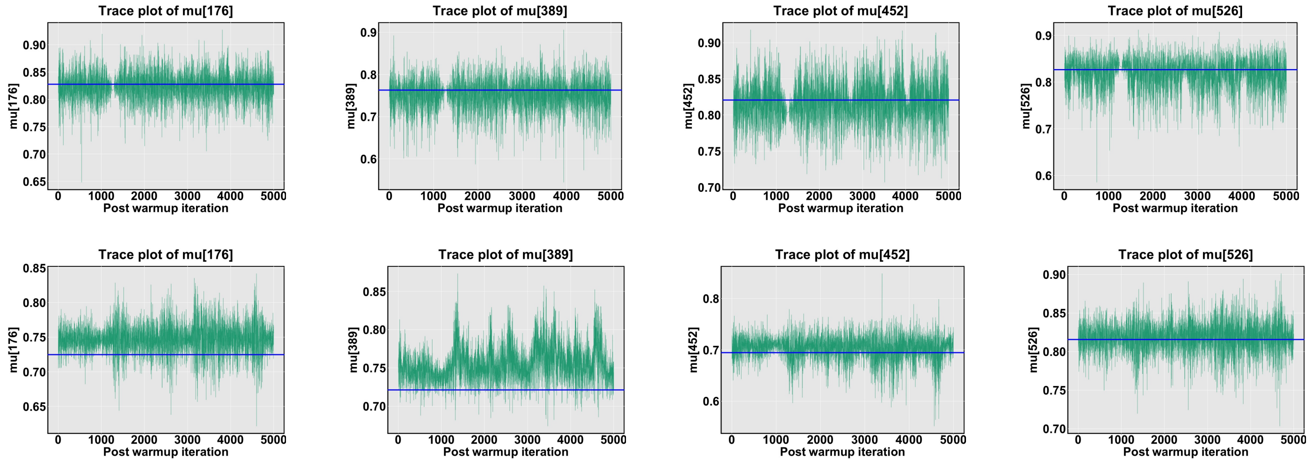


Figure S1: *Diagnostic plots with $n = 100$ (top row) and $n = 500$ (bottom row) of 4 randomly selected coordinates of the mean parameter. The blue horizontal line is the true value of $\mu$ at that coordinate.*

### S2 Selection of $M$

We vary $M \in \{5, \ldots, 30\}$ for model BR-MSIM($\mu_{it}, \psi_i$) and BR-MSIM($\mu_{it}, \psi$) and calculate BIC for each choice of $M$. Figure (S2a) suggests that for any value of M model 4 has lower BIC value than model 1 but it is difficult to observe the value of M corresponding to the lowest BIC. It is evident from figure (S2b) that BIC is lowest at $M = 22$ for the model BR-MSIM($\mu_{it}, \psi$).

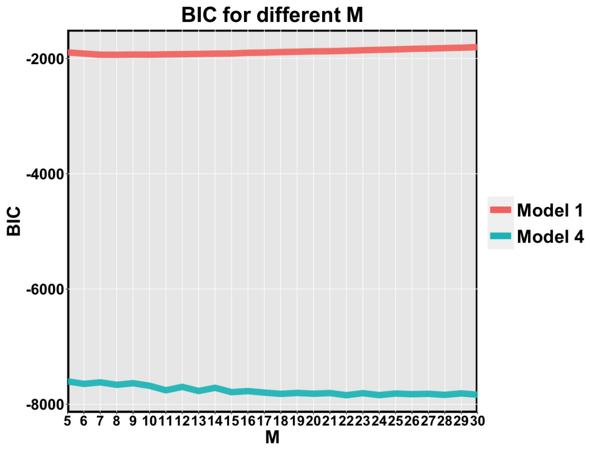


(a)

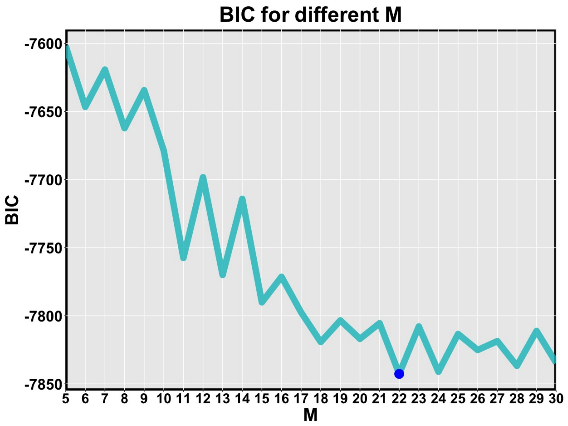


(b)

Figure S2: *In the left panel, we have the BIC values from BR-MSIM($\mu_{it}, \psi_i$) and BR-MSIM($\mu_{it}, \psi$) model for $M \in \{5, \ldots, 30\}$. The right panel shows the BIC values only from BR-MSIM($\mu_{it}, \psi$) model and the blue point denotes the lowest BIC value at $M = 22$.*

### S3 Boxplots of Euclidean distances

Here, we plot the discrepancy between the *true* and *estimated* single index, for increasing $n$.

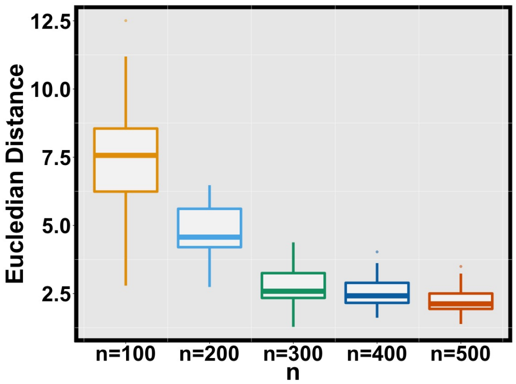


(a)

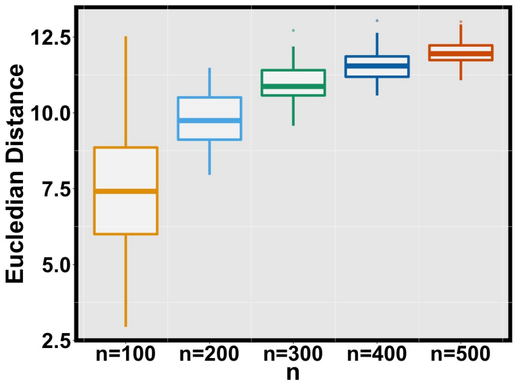


(b)

Figure S3: *Boxplot of* $100 \times d_s$ *between true and estimated single index parameters across* $50$ *replicates with increasing* $n$*. Left panel represents Simulation setting 1; right panel represents Simulation setting 2*
